# Consistent Force Field Captures Homolog Resolved HP1 Phase Separation

**DOI:** 10.1101/2021.01.06.425600

**Authors:** Andrew P. Latham, Bin Zhang

## Abstract

Many proteins have been shown to function via liquid-liquid phase separation. Computational modeling could offer much needed structural details of protein condensates and reveal the set of molecular interactions that dictate their stability. However, the presence of both ordered and disordered domains in these proteins places a high demand on the model accuracy. Here, we present an algorithm to derive a coarse-grained force field, MOFF, that can model both ordered and disordered proteins with consistent accuracy. It combines maximum entropy biasing, least-squares fitting, and basic principles of energy landscape theory to ensure that MOFF recreates experimental radii of gyration while predicting the folded structures for globular proteins with lower energy. The theta temperature determined from MOFF separates ordered and disordered proteins at 300 K and exhibits a strikingly linear relationship with amino acid sequence composition. We further applied MOFF to study the phase behavior of HP1, an essential protein for posttranslational modification and spatial organization of chromatin. The force field successfully resolved the structural difference of two HP1 homologs, despite their high sequence similarity. We carried out large scale simulations with hundreds of proteins to determine the critical temperature of phase separation and uncover multivalent interactions that stabilize higher-order assemblies. In all, our work makes significant methodological strides to connect theories of ordered and disordered proteins and provides a powerful tool for studying liquid-liquid phase separation with near-atomistic details.

## INTRODUCTION

Many proteins encoded by eukaryotic genomes contain disordered regions that do not adopt well-defined tertiary structures. ^1–6^ Disordered domains could facilitate the target search process while retaining protein-binding specificity via the folding-upon-binding mechanism. ^7,8^ It has recently become widely appreciated that another important property of these intrinsically disordered proteins (IDPs) lies in their collective behavior.^9,10^ The multivalent interactions that are intrinsic to them can drive the formation of membraneless organelles, including stress granules,^11^ P granules,^12^ superenchancers,^13^ and heterochromatin^14,15^ through the liquid-liquid phase separation mechanism. The increased protein concentration in these organelles could lead to more efficient biochemical reactions.^16,17^ Characterizing the structural details of these condensates could provide crucial insight into the function of the cellular processes. While progress is being made, connecting the atomistic properties of IDPs to the global structure, composition, and dynamics of the organelles remains challenging. ^18,19^

One prominent example of IDPs is heterochromatin protein 1 (HP1), a key component of *constitutive heterochromatin*.^20–22^ HP1 consists of ordered, conserved chromo (CD) and chromoshadow (CSD) domains connected via a variable disordered hinge region. The CD helps to recruit the protein to chromatin segments marked with histone H3 trimethylation (H3K9me3), while the CSD domain enables dimerization and also serves as a docking site for other nuclear proteins.^23^ In contrast to the canonical view that HP1 proteins are merely accessory players with no active role in chromatin organization, several studies recently found them spontaneously form phase-separated liquid droplets.^14,15,24,25^ These droplets could support new forms of chromatin structures that differ dramatically from the regular fibril conformations. ^26,27^ One noteworthy feature of HP1, which is shared by many of the proteins involved in forming membraneless compartments,^28,29^ is the presence of both ordered and disordered domains. This feature has made a high-resolution structural characterization of the full-length protein and the functional state with many proteins challenging.

Computational modeling could offer much needed structural details of IDPs and their aggregates and reveal the set of molecular interactions that dictate the stability of liquid droplets.^30^ However, the presence of both ordered and disordered domains in these proteins places a strong demand on the model’s accuracy. All-atom force fields, with their ever-improving accuracy,^31–36^ can in principle, accurately model protein conformations. They are computational expensive though, and long-timescale simulations needed to study slow conformational rearrangement and aggregation kinetics remain inaccessible for most proteins of interest. While many coarse-grained force fields have been introduced and proven effective at predicting the structures of globular proteins,^37,38^ they cannot be directly generalized to study IDPs. Separate efforts have been carried out to parameterize force fields specialized for disordered proteins.^39–42^ These force fields, while succeeding in modeling the phase separation and IDP structural heterogeneity, ^42,43^ are not advised for applications of globular proteins. Since the two classes of proteins share the same set of amino acids for their composition, it is hopeful that a unified force field can be derived to model both of them with consistent accuracy. Such a consistent force field would greatly facilitate the investigation of IDPs’ collective behaviors in large scale condensates.

In this paper, we introduce a new algorithm to parameterize coarse-grained protein force fields with implicit solvation. We generalize the maximum entropy optimization algorithm by ensuring that for globular proteins, the force field predicts an energy gap between the native conformation and the unfolded or partially folded structures. The maximum entropy optimization algorithm was developed for parameterizing transferable IDP force fields using biasing energies derived from experimental constraints. ^44,45^ Energy gap maximization, on the other hand, has been a successful strategy for deriving force fields of folded proteins.^46–49^ The resulting force field, MOFF, indeed provides a more balanced set of interactions that can predict the radii of gyration of both ordered and disordered proteins. The theta temperatures determined using MOFF classify the two types of proteins by the biological temperature at 300 K. They exhibit a striking linear relationship on the protein sequence composition. We applied MOFF to characterize the two homologs of human HP1, HP1*α* and HP1*β*. Simulations succeeded at predicting the relative size of the two proteins despite their high sequence similarity, and revealed multivalent, charged interactions that stabilize the more collapsed HP1*α* conformations. The computational efficiency of the force field enabled direct simulations of the phase separation process. These large scale simulations helped quantify the critical temperature of the two proteins and uncovered higher-order protein clusters mediated via the same interactions that cause the collapse of dimers. We anticipate MOFF to be a useful tool for studying IDPs and provide its implementation in GROMACS (https://github.com/ZhangGroup-MITChemistry/MOFF).

## METHODS

### Coarse-grained Protein Force Field

We modeled proteins with a coarse-grained representation for efficient conformational sampling and large scale simulations of phase separation. Each amino acid is represented with one bead at the *α*-carbon position. The chemical properties of each bead are provided in Table S1. The potential energy for quantifying the stability of a protein structure ***r*** is defined as

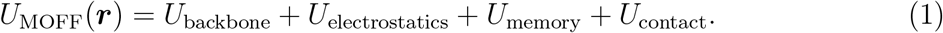

The first three terms apply to all proteins. *U*_backbone_ is responsible for maintaining the backbone geometry of the protein and consists of the bond, angle, and dihedral potentials. *U*_electrostatics_ describes electrostatic interactions between charged residues with the DebyeHückle theory. We used a distance-dependent dielectric constant to more accurately capture the change in the solvation environment upon protein folding^50,51^ (Figure S1). *U*_memory_ was applied to help stabilize protein secondary structures during force field optimization and enforce tertiary contacts in HP1 dimers. The last term *U*_contact_ is the amino acid type dependent pairwise contact potential. Parameters that quantify the strength of such interactions will be derived using the algorithm introduced in the next section to ensure consistency for both ordered and disordered proteins. The full functional form of the potential energy is given in the Supporting Information (SI).

### Amino Acid Contact Potential from Maximum Entropy Optimization

Efficient modeling of protein molecules is a long-standing interest in computational biophysics. Numerous algorithms have been introduced to parameterize coarse-grained force fields with implicit solvation for globular proteins.^37,52–57^ One class of algorithms that is of relevance to this study is inspired by the energy landscape theory,^58–61^ which states that an energy gap between the folded native structure and unfolded conformations is necessary and sufficient for ensuring reliable protein folding on a reasonable timescale. Several force fields have been introduced via maximization of the energy gap and successfully applied to predict protein structures and study folding kinetics. ^46–49^

The force fields designed for globular proteins often need to be adjusted when applied for IDPs. ^40–42^ We recently showed that maximum entropy optimization serves as an efficient means to improve the accuracy of existing coarse-grained protein force fields with experimental data.^62^ Maximum entropy optimization has become widely popular due to its relation to statistical mechanics and information theory.^63–69^ We further introduced an iterative algorithm based on maximum entropy optimization to create a transferable force field capable of reproducing the radius of gyration for various IDPs (MOFF-IDP).^44^ As detailed in the SI, the essence of this algorithm is to reparameterize the protein-specific linear bias derived from maximum entropy optimization with a transferable contact potential between pairs of amino acids,

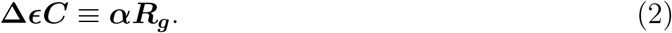

***C*** is the contact matrix that indicates the number of contacts for each pair of amino acids in a given structure. **Δ ϵ** is the list of changes to the pairwise amino acid specific contact energy. ***αR***_***g***_ is the biasing energy derived from maximum entropy optimization. When solved over an ensemble of structures collected for a large set of proteins in the training set, the changes to the contact energy matrix (**Δ ϵ**) can be determined.

It is worth noting that IDP force fields, either modified from existing ones designed for globular proteins or parameterized from scratch, often cannot be applied to predict the size and structure of folded proteins. Therefore, there appears to be a disconnect between folded and unfolded proteins. Force fields often only work on one of them, but not the other. This paper introduces a new algorithm to parameterize force fields that can be applied to model both ordered and disordered proteins with consistent accuracy.

The starting point of the algorithm is still the iterative maximum entropy optimization (see Figure 1). While our previous study focused on IDPs, here we added globular proteins as well in the training set to build a consistent force field. In addition, we borrowed ideas from the energy landscape theory to enforce that the total contact energy of the PDB structure is lower than that of any structure sampled in computer simulations, up to some tolerance. Mathematically, this constraint can be expressed as

**Figure 1:**
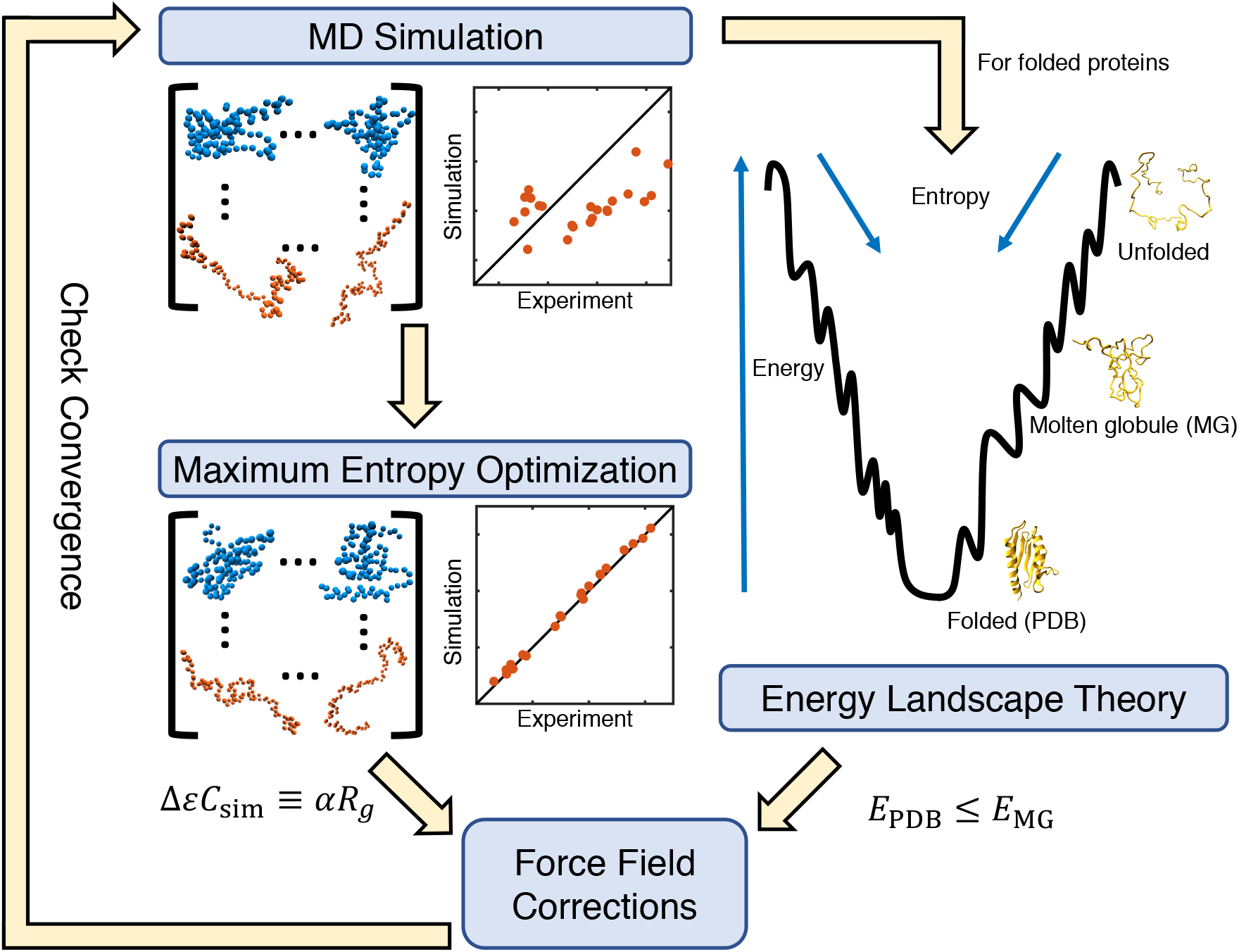
Illustration of the algorithm that combines maximum entropy optimization and energy gap constraint for force field parameterization.

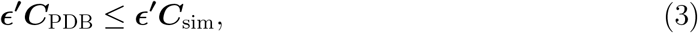

where ϵ^***′***^ = **Δ ϵ** + ϵ is the new pair-wise contact energy updated with the correction term from Eq. 2. Thus, using the interior-point algorithm, ^70^ we simultaneously solved Eq. 2 for all proteins in our training set under the constraint of Eq. 3 for the ordered proteins (Figure 1). In practice, we added an additional term, *γσ*_**sim**_ to the right hand side of Eq. 3. *γ* is a flexible tolerance parameter used to tune the strength of the constraint, and *σ*_**sim**_ is the standard deviation of the contact energy estimated for each protein using simulations performed in the previous iteration. To aid the convergence of the algorithm, we used single value decomposition (SVD) to reduce noise and placed an additional constraint on the change in amino acid contact energies from one iteration to the next. As seen previously,^44^ the relationship between energy and contact formation is not perfectly linear, requiring this entire algorithm to be done iteratively.

### Details on Molecular Dynamics Simulations

The simulations used for force field optimization were initialized from PDB structures for ordered sequences, or I-TASSER predictions for disordered sequences. ^71^ GROMACS was used to perform simulations,^72^ and the PLUMED plugin was used to incorporate biasing potentials.^73^ Proteins were placed in cubic boxes with side lengths of 50 nm to prevent them from contacting the periodic images. Each simulation trajectory consisted of six replicas at 300, 320, 340, 360, 380, and 400K, with swaps attempted every 100 steps. Langevin dynamics were used to control the temperature with a coupling constant of 1 ps, and lasted for 4 × 10^7^ steps with a time step of 10 fs. We excluded the first 10^7^ steps for equilibration and sampled protein configurations every 20000 steps. This resulted in 1500 structures per simulation, and the two trajectories combined produced 3000 structures for each protein. These structures were used to solve Eq. 2 and Eq. 3. More simulation details can be found in the SI *Section: Simulation Details on Force Field Optimization*.

Initial structures for HP1 dimers were built using RaptorX, ^74^ with the crystal structure of the HP1*α* CSD domain as a template (PDB: 3I3C). These structures were placed in cubic boxes with side lengths of 500 nm for HP1 dimer simulations. Each simulation trajectory consisted of six replicas at 300, 320, 340, 360, 380, and 400 K, with swaps attempted every 100 steps. Langevin dynamics were used to control the temperature with a coupling constant of 1 ps, and lasted for 5×10^8^ steps with a time step of 10 fs. The first 10^8^ steps were excluded for equilibration, and samples were taken every 5 × 10^4^ steps. Data from five sets of replica-exchange simulations were concatenated and analyzed, and changes from one simulation to another were statistically insignificant. Clustering was done following the gromos clustering algorithm.^75^ More details on the dimer simulations are available in the SI *Section: HP1 Dimer Simulation Details*. Slab simulations performed to study HP1 phase separation are explained in the SI *Section: HP1 Slab Simulation Details*.

## RESULTS AND DISCUSSION

### Parameterization of the Consistent Force Field

We applied an iterative optimization algorithm to parameterize the tertiary contact potentials of a coarse-grained force field (Eq. 1) and ensure consistent accuracy for both IDPs and ordered proteins. Details of the algorithm can be found in the *Methods Section*. As illustrated in Figure 1, it matches the simulated radius of gyration of protein molecules with experimental values via maximum entropy optimization. ^62^ In addition, we enforce the constraint that, for folded proteins, the energy of the native structure is lowest, i.e., there is a gap between folded and unfolded configurations. This gap is necessary to ensure reliable protein folding into the native state. A total of 23 proteins, including seven ordered and 16 disordered (Table S2), were included in the training set. We tracked the percent error given by

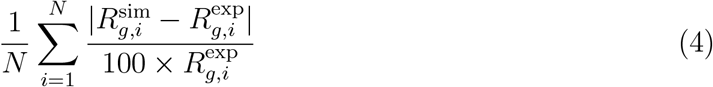

to measure the improvement caused by our optimization. The sum is taken over all *N* = 23 proteins in our training set, and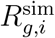 and 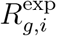 are the simulated and experimental radius of gyration respectively. In addition to tracking the percent error on our training set, we monitored the performance of the force field on an independent validation set as well (Table S3). The validation set includes four ordered and four disordered proteins. We terminated the optimization when the percent error on the validation set fails to decrease upon two consecutive iterations. Starting values of the amino acid contact energies were set as the Miyazawa-Jernigan (MJ) potential^76^ scaled by a factor of 0.4. The scale factor was determined as the value with the least percent error for proteins in the training set (Figure S2). We gradually relaxed the constraint defined in Eq. 3 by increasing the tolerance term along the iterations (Figure S3).

As shown in Figure S3, the iterative algorithm succeeds in gradually improving the force field accuracy. The percent error begins at 33.6, but reaches 9.6 by the final iteration. Importantly, the resulting force field (MOFF) outperforms the initial one built from the MJ potential for every protein in the training set (Figure 2A). When examining the validation set, we again noticed that (MOFF) improves relative to both MJ and MOFF-IDP, where MOFF-IDP is a force field parameterized specifically for IDPs using the maximum entropy optimization algorithm. ^44^ The percent error for proteins in the validation set is 17.9, compared to 51.2 and 41.2 for MOFF-IDP and MJ respectively (Figure 2B). In particular, we note that the new force field significantly improves on the ordered proteins (low *R*_*g*_) relative to MOFF-IDP. This improvement likely stems from the newly added ordered proteins in the training set for force field parameterization. MOFF performs better than the other two force fields on all but one protein, 4cpv. The poor performance may be due to the presence of a calcium ion in the PDB structure, which was absent in our model due to the use of implicit solvation.

**Figure 2:**
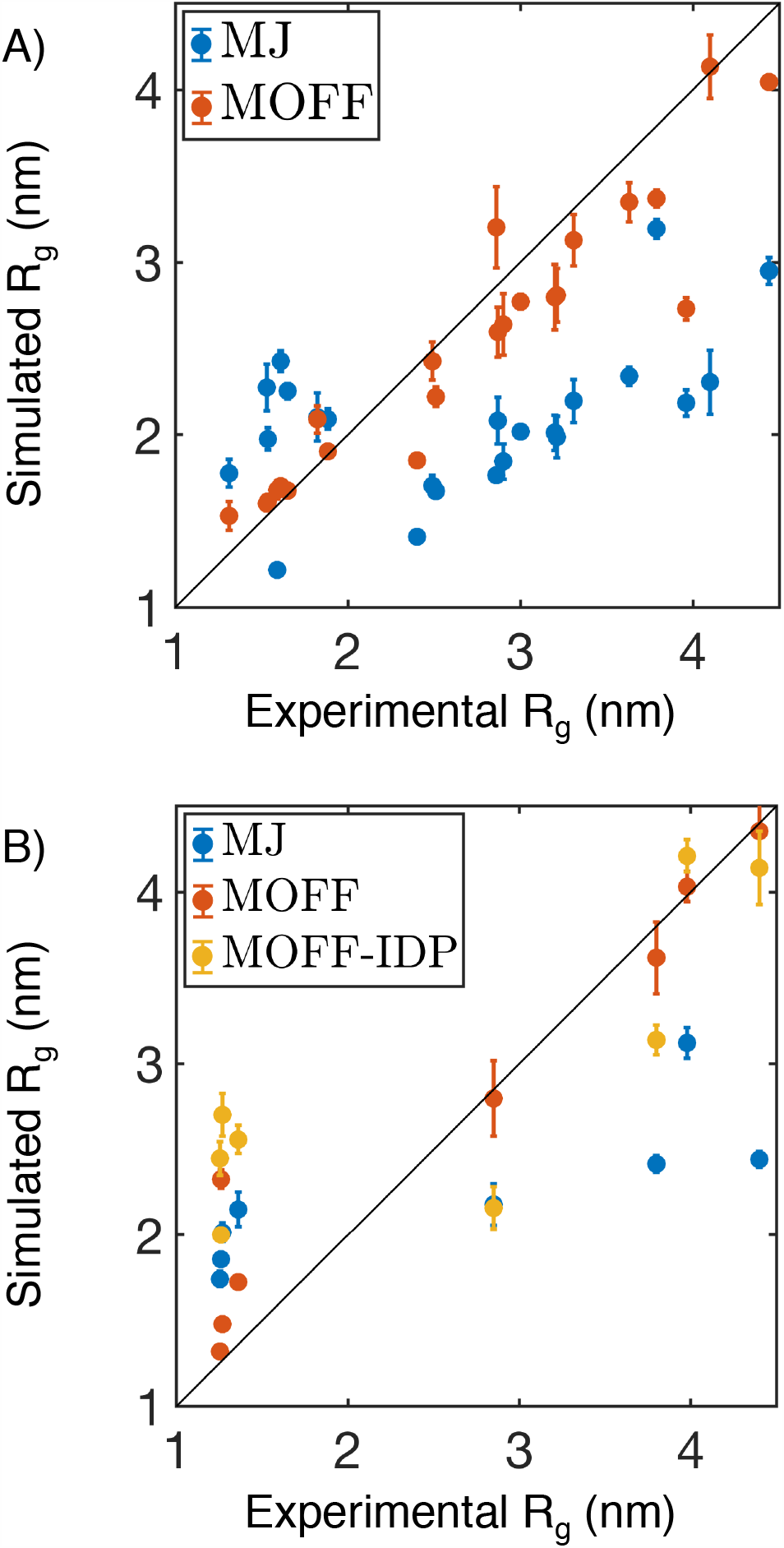
Comparison between experimental and simulated radius of gyration (*R*_*g*_) for protein molecules in the training (A) and validation (B) set. In addition to the force field introduced in this paper (MOFF, red), we included simulation results using the Miyazawa-Jernigan potential (MJ, orange) and a previous version of MOFF optimized for IDPs (MOFF-IDP, blue) as well. Error bars represent standard deviation after block averaging.

The optimization also succeeded at ensuring that the folded structure is lower in energy than the unfolded configurations. The energy gap, defined as the difference between the mean contact energy of the simulated structures and the contact energy of the folded structure (Figure 3A), is indeed positive for all the globular proteins in the training set (Figure 3B). Upon a close inspection of simulated configurations, we observed deviations from the native conformation, with the probability distribution of the root mean squared displacement (RMSD) from the PDB structure peaking around 1.5 nm (Figure S4). It is worth noting that several proteins do sample configurations close to the native state with RMSD less than 0.5 nm. In particular, 1wla shows a bimodal distribution with one of the peaks located at 0.5 nm (Figure 3C). Furthermore, simulated annealing simulations were able to predict structures with small RMSD values for three of the seven proteins (Figure S5). These improvements seem to be related to secondary structure content. For example, MOFF performs best on 1wla, which is an *α*-helical protein, while it performs worst on 5tvz, which is a *β*-sheet protein. More refined models that explicitly consider side chains are probably needed to ensure more consistent performance in structural prediction for all globular proteins.

**Figure 3:**
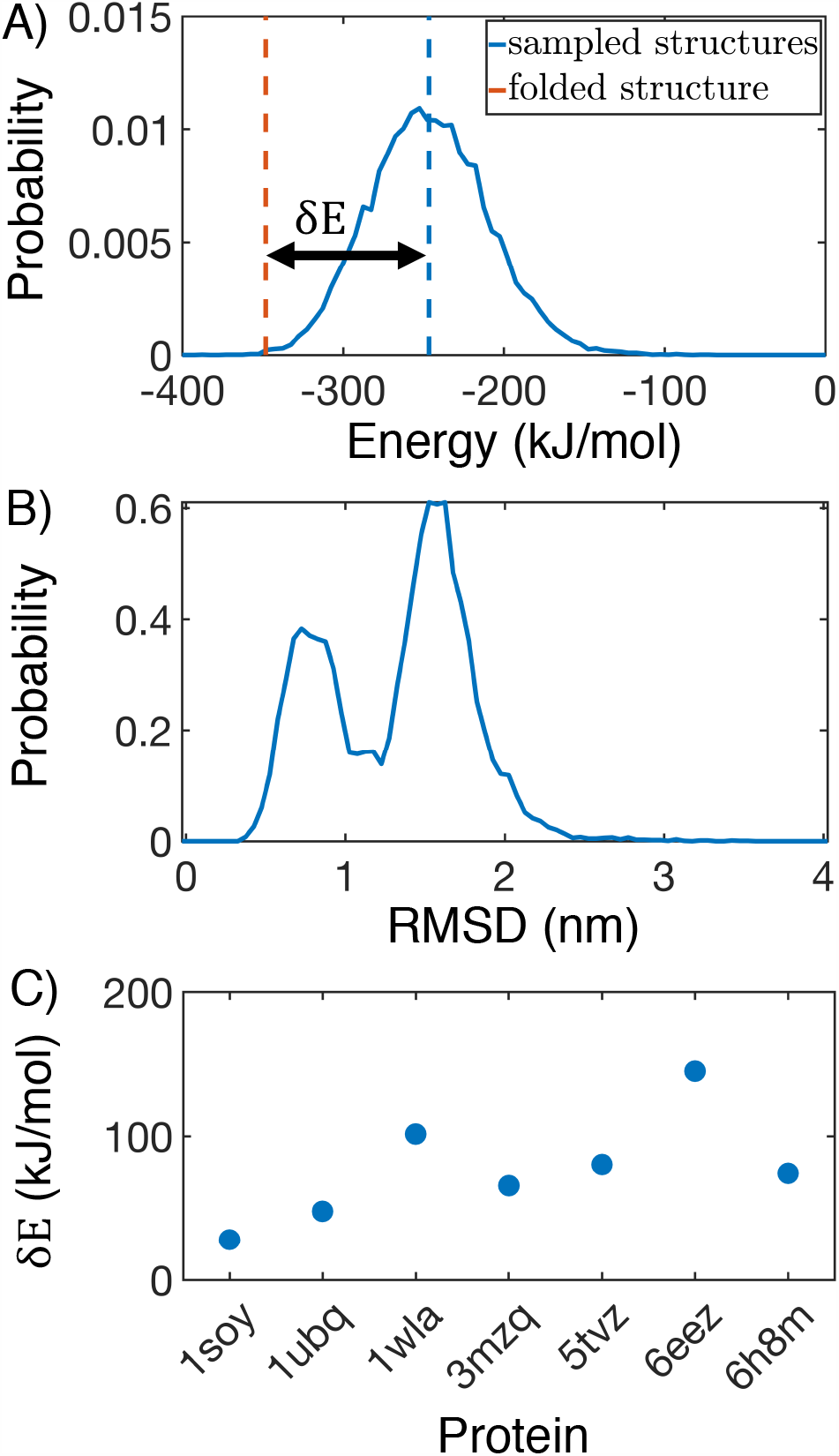
MOFF succeeds in creating energy gaps for folded proteins. (A) Contact energy of the folded structure (orange) relative to those sampled in simulation (blue) for 1wla. The energy gap *δ E* is defined as the difference between the folded energy and the average simulated energy. (B) Probability distribution of the root mean squared displacement (RMSD) relative to the folded structure for 1wla at 300 K. (C) *δ E* for all folded proteins in the training set.

### MOFF Identifies Sequence Features to Differentiate Ordered and Disordered Proteins

To gain insight into the molecular interactions that dictate MOFF’s success in predicting protein sizes, we performed a hierarchical clustering on the contact energy matrix based on euclidean distances between column vectors. The resulting clusters generally sort the amino acids into groups with increasing hydrophobicity (Figure 4A and Table 1). When evaluating the frequency of encountering the various amino acid clusters in the ordered and disordered proteins from our training set, we found that cluster 3 consists of mainly hydrophilic amino acids and is significantly enriched in disordered proteins. On the other hand, the hydrophobic cluster 6 is more often seen in folded proteins. Similar trends have been seen in more expansive studies, which compared the frequency of amino acids in the DisProt database with the frequency in the PDB.^77,78^ The example configurations shown in Figure 4C further highlights the spatial distribution of the various clusters in an ordered and a disordered protein.

**Figure 4:**
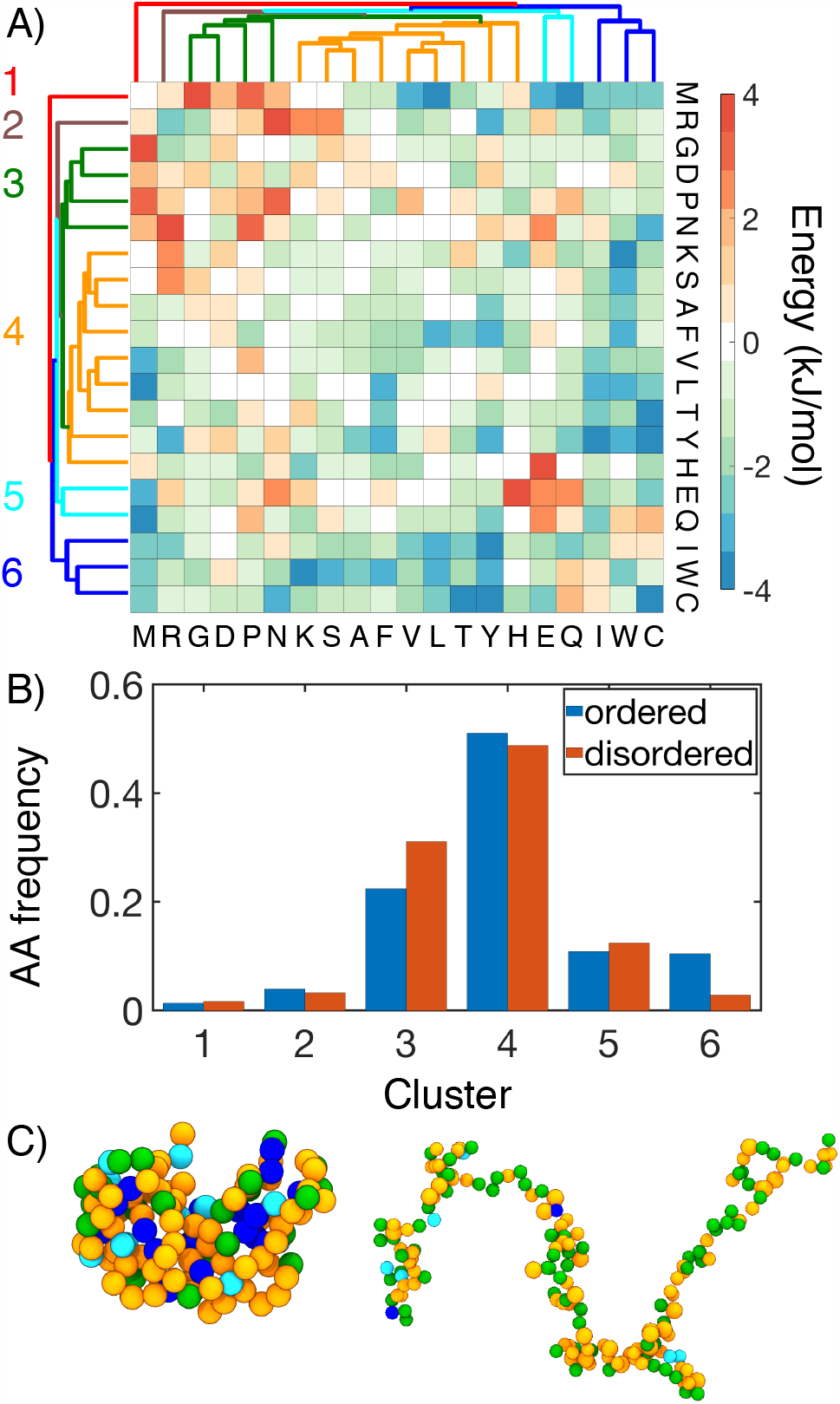
MOFF differentiates “ordered” and “disordered” amino acids with distinct contact energy patterns. (A) Hierarchical clustering of MOFF contact energy between amino acid pairs (ϵ_*IJ*_ in Eq. S16). (B) Comparison of the amino acid frequency by cluster in the ordered (blue) and disordered (orange) portions of our training set. (C) Spatial distribution of amino acid clusters in example structures of an ordered (3mzq, left) disordered protein (NSP, right). Amino acids are shown in the same coloring scheme for clusters as in part A.

**Table 1:** Amino acid clusters determined from hierarchical clustering of MOFF contact energy.

| Cluster | Amino Acids |
| --- | --- |
| Cluster 1 (red) | MET |
| Cluster 2 (brown) | ARG |
| Cluster 3 (green) | PRO ASN ASP GLY |
| Cluster 4 (orange) | HIS PHE LYS SER ALA VAL LEU THR TYR |
| Cluster 5 (cyan) | GLN GLU |
| Cluster 6 (blue) | ILE TRP CYS |

The presence of well-defined amino acid clusters that sets apart ordered proteins from disordered ones provides an intuitive explanation of the distinct size distribution of the two protein types. The “disordered” amino acids from cluster 3 are mainly repulsive and will lead to the more expanded structure seen in IDPs; the “ordered” amino acids from cluster 6, on the other hand, are attracted to most of the other residues and tend to localize in the interior of collapsed proteins. The interaction energy among amino acids also explains the performance of the three force fields shown in Figure 2. Compared to the MJ potential (Figure S6), interactions between hydrophilic residues are more repulsive in MOFF. Similar results were seen with MOFF-IDP, where repulsive interactions rescue IDPs from over collapse. However, when ordered proteins are also included in the training set, we see these repulsive interactions grow stronger, as well as the development of stronger contact energy among hydrophobic residues. These changes are likely necessary to balance the collapse of ordered sequences and the swelling of disordered sequences.

We next attempt to establish a more quantitative relationship between protein size, interaction energy, and the sequence. Towards that end, we first compute the theta temperature, *T*_*θ*_, by monitoring the change of scaling exponent *ν* as a function of temperature. *ν* measures the variation of spatial distance between two residues *i* and *j* versus their linear distance in sequence, *R∝* |*i − j* |. At *T*_*θ*_, proteins behave like an ideal Gaussian chain and *ν* = 1*/*2. As the temperature decreases from above to below *T*_*θ*_, one expects the proteins to become more compact and transition from swollen polymers to collapsed globules, as can be seen in Figure 5A-B. *T*_*θ*_, therefore, provides a direct measure of the interaction strength within a protein. Strikingly, MOFF predicted that the biological temperature provides a clear cut between ordered and disordered proteins, with the corresponding *T*_*θ*_ above and below 300 K, respectively. Such a partition indicates that MOFF treats the two types of proteins as polymers in a poor or good solvent and is consistent with the results shown in Figure 2. The interaction potential of a protein and *T*_*θ*_ is ultimately dictated by the underlying sequence. Given the similarity among amino acids in their contact energy, we wondered whether a simple relationship between *T*_*θ*_ and the sequence composition can be found. Using the least absolute shrinkage and selection operator (LASSO), ^79^ we determined a linear regression model to fit *T*_*θ*_ with a minimal set of amino acid clusters identified in Figure 4. The model that minimizes the mean squared error adopts the following expression

**Figure 5:**
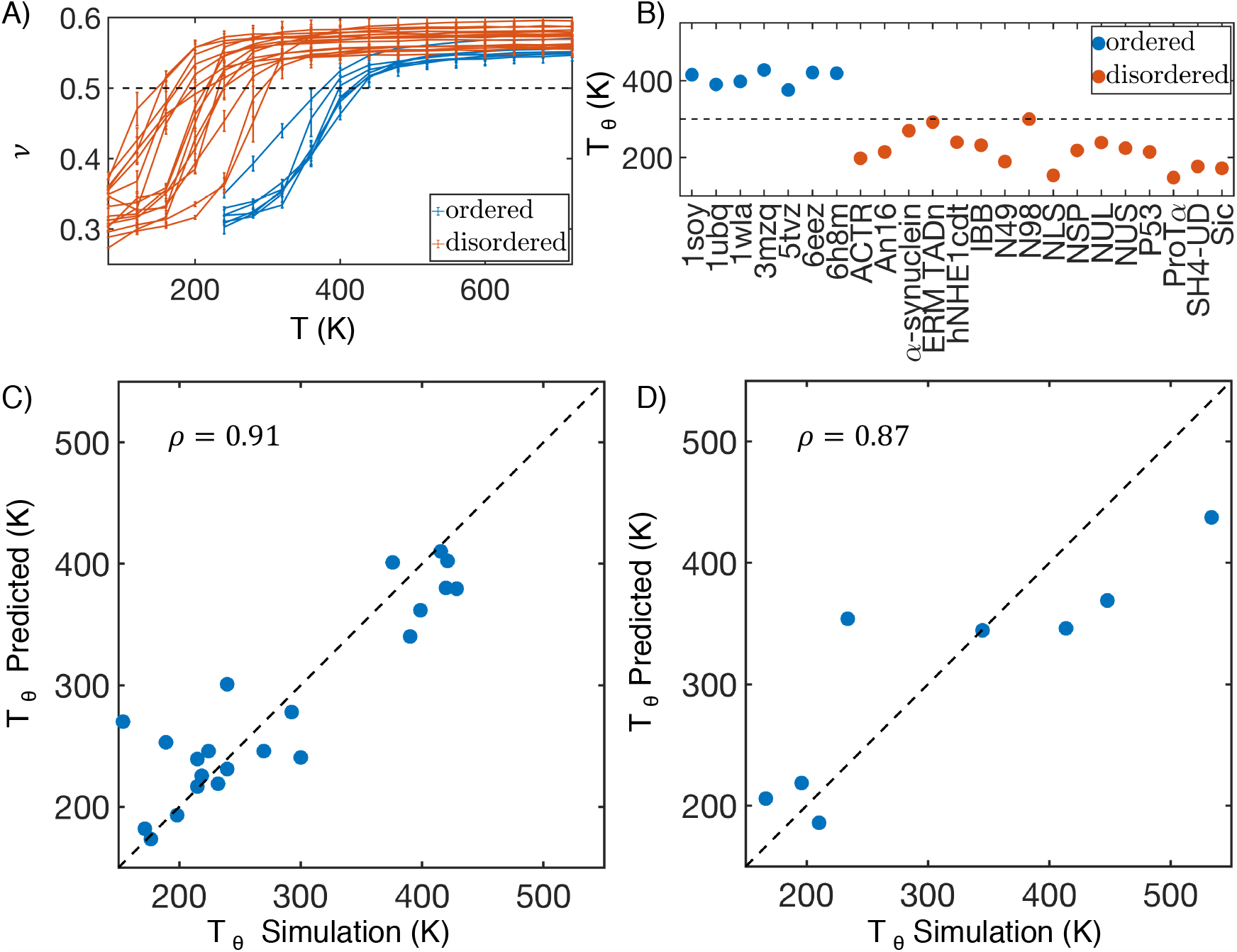
MOFF uncovers a linear relationship between protein theta temperature and sequence composition. (A) Polymer scaling exponent (*ν*) as a function of *T* for ordered (blue) and disordered (orange) protein sequences in our training set. Error bars represent standard deviation after block averaging. (B) Theta temperature (*T*_*θ*_) for proteins in the training set. The 300K mark is highlighted as a guide for the eye. (C, D) Comparison between values of *T*_*θ*_ predicted using a linear equation of sequence composition (Eq. 5) and determined from molecular dynamics simulations for proteins in the training (C) and validation (D) set. *ρ* is the Pearson correlation coefficient between the two data sets, and the dashed black line represents perfect agreement.

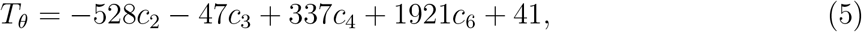

where *c*_*i*_ is the percent of the sequence that is from cluster *i* (Figure S7). The fitted values are strongly correlated with the simulated ones despite our neglect of clusters 1 and 5 in the expression (Figure 5C). This expression supports the notion of hydrophobic effect ^80,81^ since amino acids from clusters 4 and 6 are mostly hydrophobic, and they contribute positively to the theta temperature by promoting protein collapse. It is also consistent with our conclusion that IDPs are more expanded because of their enrichment in residues from cluster 3. Arg, which is the sole residue in cluster 2, appears to be effective at reducing *T*_*θ*_ as well, potentially resulting from electrostatic repulsion.

We further computed *T*_*θ*_ for proteins in the test set and observed separation between ordered and disordered proteins at 300 K again (Figure S8). Notably, *T*_*θ*_ computed from replica exchange simulations are in good agreement with the values predicted using Eq. 5, with a Pearson correlation coefficient of 0.87 between the two (Figure 5D). Therefore, the linear relationship between *T*_*θ*_ and the sequence composition is general and transferable.

### Multivalent Interactions Differentiate HP1 Homologs

After validating its accuracy in modeling the size of both ordered and disordered proteins, we applied MOFF to characterize the structure and phase behavior of human HP1. We first investigated the difference between the two isoforms, HP1*α* and HP1*β*. Previous experimental studies on the HP1 dimers showed that HP1*β* takes an open conformation, while HP1*α* is more collapsed.^14,82^ The size difference is particularly striking considering the sequence similarity between the two proteins. As shown in Figure 6A, the ordered regions that include chromodomain and chromoshadow domain share a sequence identity over 80%. Even the disordered N-terminal extension, hinge region, and C-terminal extension have a sequence similarity of over 30%. ^83^

**Figure 6:**
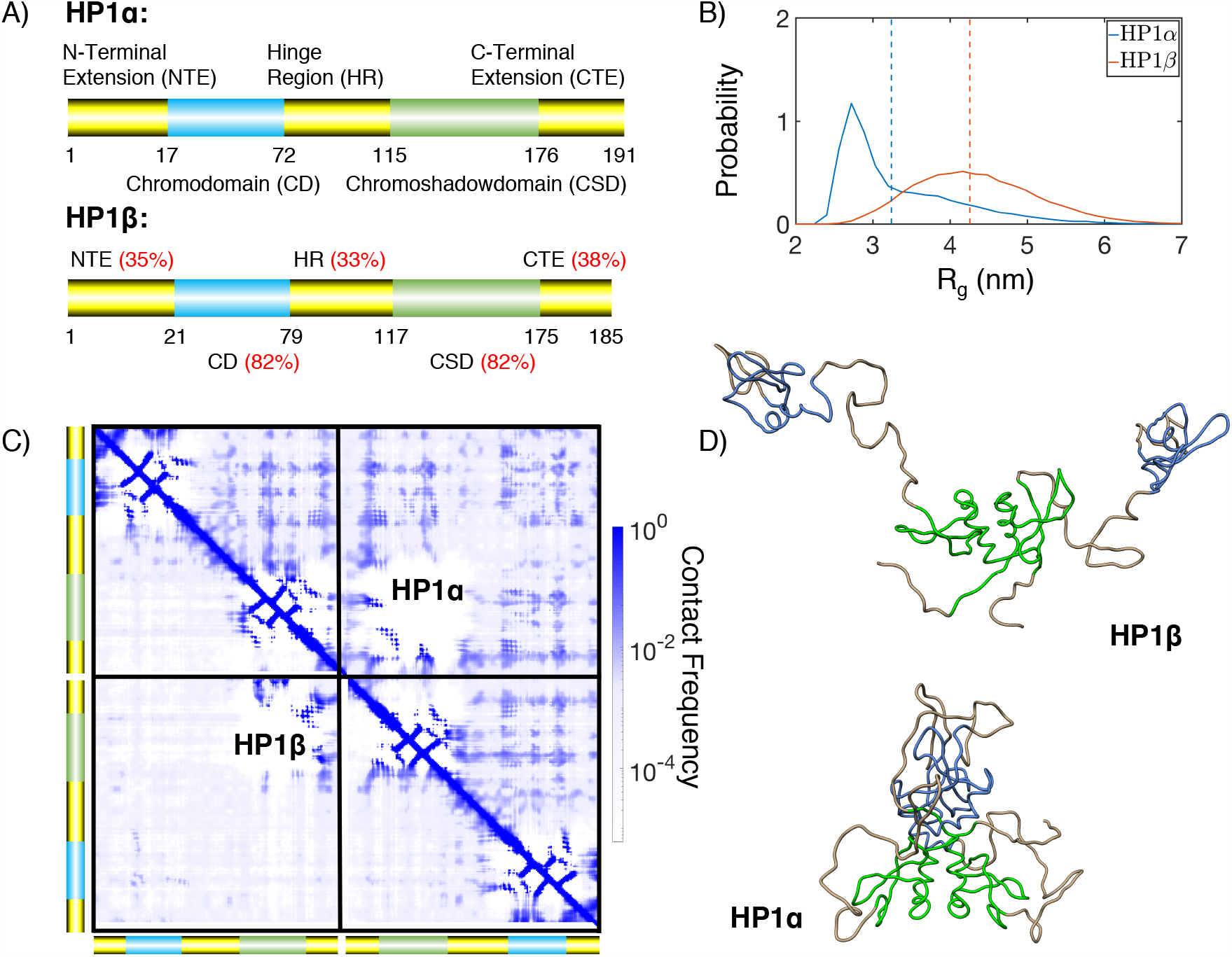
MOFF resolves the structural difference between HP1*α* and HP1*β*. (A) Cartoon diagrams for the two HP1 homologs, with the disordered regions shown in yellow and ordered regions in blue and green. The red numbers indicate sequence identity between the two proteins for various regions.^83^ (B) Probability distributions of the radius of gyration (*R*_*g*_) for HP1*α* (blue) and HP1*β* (orange). Dashed lines show mean values of each distribution. (C) Contact maps of HP1*α* (top right) and HP1*β* (bottom left), with cross-dimer interactions shown in the diagonal quadrants. (D) Representative structures of HP1*β* and HP1*α* determined from the most populated cluster. The coloring scheme is the same as in part A.

We performed replica exchange simulations of the HP1 dimers. Similar to experimental studies, we find that HP1*α* takes a more collapsed configuration at 300 K, with *R*_*g*_ = 3.24 ± 0.08 nm, compared to *R*_*g*_ = 4.26 ± 0.08 nm for HP1*β* (Figure 6B). These results compare favorably to experimental values of 3.59 nm and 4.7 nm, respectively.^14,82^

We found that multivalent interactions between charged residues cause the more compact configurations of HP1*α*. These interactions are evident in the contact maps shown in Figure 6C for HP1*α* (upper triangle) and HP1*β* (lower triangle). The off-diagonal quadrants correspond to interactions between the two monomers. For HP1*β*, these interactions are mostly limited to the dimerization of the chromoshadow domain. The contacts are more widespread in HP1*α* and arise from charged interactions between positive residues from the chromodomain or hinge regions and the negative counterparts of the chromoshadow or C-terminal extension domains. Such contacts can be readily seen in the central structure of the most populated cluster shown in Figure 6D. The clustering was performed over the simulated structural ensemble based on RMSD (Figure S9). On the contrary, we see that in HP1*β*, the hinges are completely extended, and there are minimal interactions between the two monomers.

### Homolog Specific HP1 Phase Separation

The multivalent interactions found in HP1*α* are weak and can come undone to form more extended structures. These structures give rise to the long tail of the probability distribution of *R*_*g*_ (Figure 6B) and become more stable at higher temperatures (Figure S10). The extended configurations could facilitate contacts between different dimers to promote large-scale protein aggregation and phase separation.

We performed slab simulations^40,43^ with 100 dimers of HP1*α* and HP1*β* to directly probe their phase behavior. Starting from configurations with a high concentration of protein molecules in a cubic box, we extended the simulation box along the *z* direction by 20 times. The system was then relaxed under constant volume simulations with periodic boundary conditions to reach desired temperatures. After equilibration, protein molecules will either disperse throughout the simulation box (Figure 7A), or stabilize into two phases with different concentrations (Figure 7B). A total of ten simulations with temperatures ranging from 150 K to 400 K were performed for each protein. We monitored the dynamics of these simulations by tracking the size of the largest cluster of HP1 dimers as a function of time. As shown in Figure S11, while low-temperature simulations preserve the initial dense phase, proteins begin to shake off as the temperature increases, leading to a drop in the cluster size. Using the identified HP1 clusters, we further partitioned the system into two phase regimes and computed the corresponding protein concentration in each phase at high temperatures (Figure 7C). We then determined the critical temperature, *T*_*C*_, by fitting the concentrations as a function of temperature using the following expression

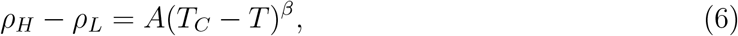

**Figure 7:**
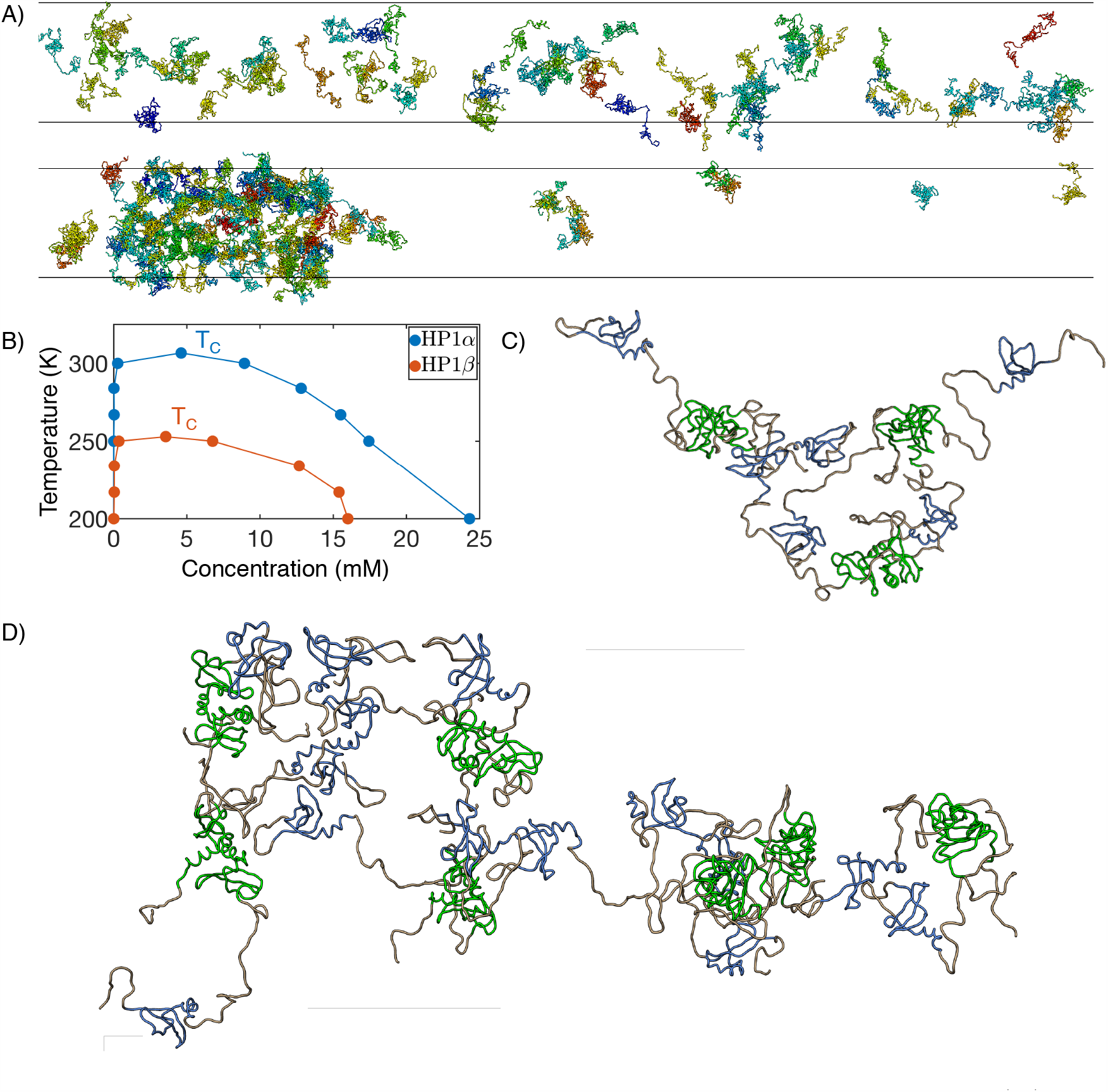
MOFF enables quantitative simulations of HP1 phase separation. (A) Example configurations from slab simulations of HP1*α* at 350 (top) and 300 (bottom) K. (B) Phase diagram and critical temperature for HP1*α* (blue) and HP1*β* (red). (C, D) Representative cluster structures formed with three (C) and seven (D) HP1*α* dimers. The coloring scheme is the same as in Figure 6A.

with *β* = 0.325.^43^ The resulting values for HP1*α* and HP1*β* are 306.7 and 252.9 K respectively (Figure S12). The lower *T*_*C*_ of HP1*β* indicates that its dense phase is less stable and will fall apart at lower temperatures than HP1*α*. This conclusion is in agreement with experimental observations that HP1*α* can phase separate *in vitro* at room temperature, while HP1*β* cannot. ^14,25^

We found that the multivalent interactions that drive the collapse of HP1*α* dimer indeed mediate inter-dimer interactions. For example, contacts between the N terminal extension (NTE) and the NTE, Chromodomain (CD), and hinge region of other dimers can be readily seen in the example clusters shown in Figure 7D,E. The interactions are also evident in the contact map between dimers (Figure S13 red box). Our results agree with previous experimental studies, which proposed that NTE bridges dimers through charged interactions.^14^ In addition, we also observed inter-dimer interactions mediated by C-terminal extension (CTE) (Figure S13 green box). The inter-dimeric contacts are largely preserved in clusters of HP1*β*, though at a weaker strength (Figure S14). These highly patterned, multivalent interactions are, therefore, consistent across homologs.

## CONCLUSIONS

In this work, we introduced an algorithm to parameterize force fields that can be used to study both ordered and disordered proteins with consistent accuracy. We combined principles of the energy landscape theory with the maximum entropy optimization to recreate experimental radii of gyration for ordered and disordered proteins while simultaneously ensuring that the folded structures are lower in energy than the unfolded ones. The resulting force field, which we term as MOFF, indeed outperforms two existing force fields that work well for globular proteins or IDPs to recreate protein size. The force field further helped identify two clusters of amino acids critical for determining protein size and the theta temperature. When applied to the human HP1, MOFF successfully resolves the structural difference between two homologs with high sequence similarity. It identified non-specific, charged interactions that stabilize a more collapsed configuration in HP1*α* than HP1*β*. In addition, the force field was shown to be computationally efficient for studying phase separation. We determined the critical temperature for the two homologs. The values agree with qualitative experimental observations. Our simulations also provided structural insight into the condensates. The multivalent interactions found in dimers can now bridge contacts across dimers to mediate cluster formation.

We note that while MOFF can reliably predict the size of globular proteins, it has not yet achieved consistent accuracy for *de novo* structure prediction. Prior studies have shown that more than one bead per amino acid is needed to fully capture side chains’ chemical complexity and accurately model their tight packing. ^38,84^ In addition, a more refined contact potential between amino acids to differentiate short-range direct contacts from long-range ones mediated by water molecules may prove beneficial.^49,85^ Accounting for these known biophysical properties of globular proteins could improve the force field’s accuracy in modeling IDPs as well. Finally, more advanced algorithms that maximize the ratio of the folding temperature versus the glass transition temperature can be adopted to better sculpt the funneled energy landscape for globular proteins. ^46^

## Acknowledgement

This work was supported by the National Institutes of Health (Grant 1R35GM133580-01). A.L. acknowledges support by the National Science Foundation Graduate Research Fellowship Program.

## Supporting Information Available

The Supporting Information is available free of charge on the ACS Publications website at DOI: XXX.

- Constrained Maximum Entropy Optimization Algorithm; Mathematical Expressions of Energy Function; Simulation Details on Force Field Optimization; Determination of *T*_*θ*_; HP1 Dimer Simulation Details; HP1 Slab Simulation Details; Distance dependent dielectric; Optimization of MJ; Optimization of MOFF; RMSD of training set at 300K; RMSD of training set from annealing; Contact energy of MJ and MOFF-IDP; LASSO fit of *T*_*θ*_; Fitting *T*_*θ*_ to test set; RMSD clustering of HP1 dimers; *R*_*g*_ of HP1 dimers; Cluster size from HP1 slab simulations; Determination of *T*_*C*_ for HP1; HP1 intra-dimer and inter-dimer contact map for HP1*α*; HP1 intra-dimer and inter-dimer contact map for HP1*β*; Example of contact potential; Fit HP1 CSD to all-atom simulations; Amino acid masses, charges, and sizes (*σ*) used in simulation; Description of proteins in the training set; Description of proteins in the validation set; Description of folded protein structures; Protein sequences used in this study.

## TOC IMAGE

**Figure 8:**
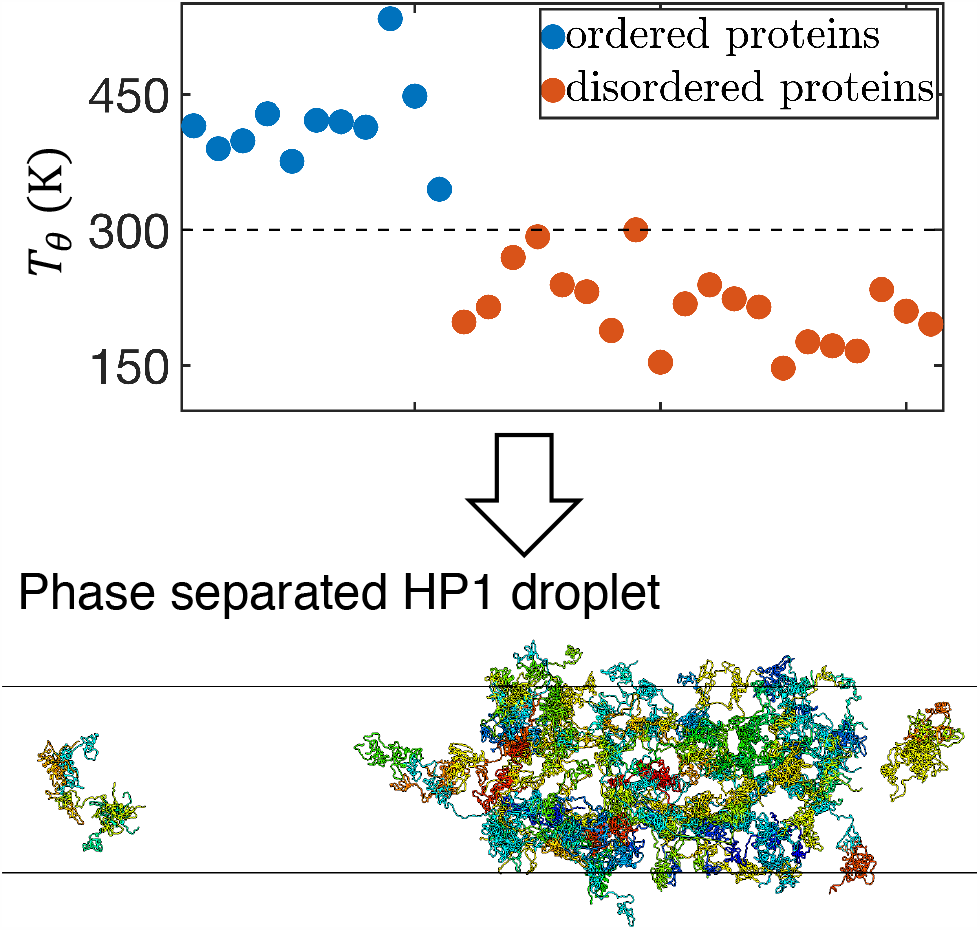
TOC Graphic.

